# Lilikoi V2.0: a deep-learning enabled, personalized pathway-based R package for diagnosis and prognosis predictions using metabolomics data

**DOI:** 10.1101/2020.07.09.195677

**Authors:** Xinying Fang, Yu Liu, Zhijie Ren, Yuheng Du, Qianhui Huang, Lana X. Garmire

**Author notes:** These authors contributed equally to the work (Xinying Fang, Yu Liu).

## Abstract

Previously we developed *Lilikoi*, a personalized pathway-based method to classify diseases using metabolomics data. Given the new trends of computation in the metabolomics field, here we report the next version of *Lilikoi* as a significant upgrade. The new *Lilikoi* v2.0 R package has implemented a deep-learning method for classification, in addition to popular machine learning methods. It also has several new modules, including the most significant addition of prognosis prediction, implemented by Cox-PH model and the deep-learning based Cox-nnet model. Additionally, *Lilikoi* v2.0 supports data preprocessing, exploratory analysis, pathway visualization and metabolite-pathway regression. In summary, *Lilikoi* v2.0 is a modern, comprehensive package to enable metabolomics analysis in R programming environment.

## INTRODUCTION

Metabolomics is an increasingly popular platform to systematically investigate metabolites as potential biomarkers for diseases (1). With the rapid development in this field, data analysis is becoming a critical component to interpret and apply the results for translational and clinical research. However, currently the majority of metabolomics analysis workflows are provided as web-apps (1), limiting its adaptation by the bioinformatics community, and/or integration with other omics workflows in a programmable matter.

To address such needs, previously we developed *Lilikoi*, a personalized pathway-based method to classify diseases using metabolomics data (2). Different from other metabolomics analysis packages, the personalized and pathway-based representation of metabolomics features is the highlight of *Lilikoi* version 1 (v1). *Lilikoi* v1 enables classifications using various machine learning methods. It has four modules: feature mapper, dimension transformer, feature selector, and classification predictor (2).

Here we report *Lilikoi* v2.0, as a significant upgrade for *Lilikoi* v1. The new mission is embarked by several recent trends or needs in the research community. Firstly, given the recent applications of deep-learning in the metabolomics and other genomics fields (3–9), it is important to enable metabolomics researchers to investigate such new approaches. We thus implemented a deep-learning neural network, as a new method in the classification module. Secondly, metabolomics have the potential to be prognosis markers (10), however, currently rarely metabolomics data analysis workflow is available for prognosis modeling and prediction. We herein implemented multiple methods for prognosis prediction, including Cox-Proportional Hazard (Cox-PH) model and *Cox-nnet*, a neural-network based model (5). Thirdly, we augmented the pathway-based metabolomics analysis with metabolite-pathway regression and pathway visualization. Last but not least, we also include additional preprocessing methods for metabolomics data analysis (eg. normalization, imputation) and tools for exploratory data analysis (eg. PCA and t-SNE analysis, and source of variation analysis). In summary, *Lilikoi* v2.0 is a more mature, comprehensive and modern package to empower the metabolomics community.

## Methods

### Datasets

Three breast cancer metabolomics datasets were used to demonstrate the new functionalities of *Lilikoi* v2.0. The first set was downloaded from the Metabolomics Workbench (https://www.metabolomicsworkbench.org/) project ID PR000284 (11), which used 207 plasma samples (126 breast cancer cases and 81 control cases) from a previous study (2). The second metabolomics dataset is from a biobank at the Pathology Department of Charité Hospital, Berlin, Germany. It contains 271 breast cancer samples, where 204 samples are ER+ and 67 samples are ER-(12). The third dataset is shared by authors from an original National Cancer Institute (NCI) study, composed of 67 breast tumor samples and 65 tumor-adjacent noncancerous tissues (10). In our analysis, we only used the 67 breast tumor samples for prognosis modeling.

### Data preprocessing

For data preprocessing, we added normalization and imputation methods. Three normalization methods (standard normalization, quantile normalization and median-fold normalization) were implemented. We used the normalize.quantiles function in the *preprocessCore* package (13) to perform the Quantile normalization. For imputation, we used k-nearest neighbors method as the default method to impute missing values. The knn imputation was done by the impute.knn in the *impute* R package (14).

### Exploratory analysis

Principal Component Analysis (PCA) is a feature selection technique (15). It extracts the most important information in high-dimensional datasets. The t-SNE plot is a dimension reduction method to help users to visualize high-dimensional data (16). We implemented the PCA and t-SNE plots in *Lilikoi v2.0* via the *M3C* package (17). We also added the source of variation analysis (SOV) for exploratory analysis, implemented by the Anova function in the *car* package (18). SOV identifies the relationships between confounders and metabolomics data, based on ANOVA tests (19, 20). Any clinical variable with F-score bigger than the error term, whose F-score is 1, is deemed a confounder.

### Metabolite to pathway level transformation

Lilikoi uses the *Pathifier* algorithm to perform the metabolites-pathway dimension transformation per sample (21). For each pathway *P* in each patient *i*, a pathway dysregulation score (PDS) *D*_*P*_*(i)* between 0 and 1 is generated, based on the metabolites associated with this pathway. A larger PDS value represents a higher degree of dysregulation (larger deviation from the normal controls). As the result of the dimension transformation, a new pathway-level matrix is constructed, which can be used to substitute the original metabolomics profile matrix, for downstream classification or prognosis modeling.

Briefly, a PDS score *D*_*P*_*(i)* is calculated as the following: in the high-dimensional space *d*_*P*_ made of metabolite vectors (where each metabolite belongs to pathway P), all samples form a data cloud, where sample *i* is a data point x_i_. The principle curve *S*_*P′*_ in this space *d*_*P*_ is then computed using Hastie and Stuetzle’s algorithm (22). For each sample, the data point x_i_ is projected onto the principle curve *S*_*P′*_. The dysregulation score *D*_*P*_*(i)* of sample *i* is then defined as the distance from the start of the principal curve to the projected point on this curve. More details of applications of *Pathifier* on biomarker studies (prognosis or diagnosis) can be found in earlier publications (2, 23, 24).

### Deep learning for classification

The deep learning algorithm in *Lilikoi v2.0* is based on the *h2o* package (25). It used a multi-layer neural network trained with stochastic gradient descent search to predict the diagnosis results. For the neural-network configuration, users are free to set parameters including activation function, hidden layer size, drop-out ratio, L1 and L2 regularization, size of one batch and adaptive learning rate decay factor. Users can also incorporate other control parameters like random discrete to optimize the hyperparameter setting to achieve the best deep learning performance.

Lilikoi *v2.0* supports users to run hyperparameter grid search on multiple deep learning models to achieve the best classification results. The activation functions are set as “Rectifier” or “Tanh”. Seven hiddenlayer configurations are pre-set for selections: 1 hidden layer setting (100 or 200 neurons), 2 hidden layer setting (10, 20 or 50 neurons for each layer), 3 hidden layers with 30 neurons for each, and 4 hidden layers with 25 neurons for each. The input dropout ratio options range from 0 to 0.9 with 0.1 increment. The number of global training samples per iteration is set to 0 or −2, where 0 means 1 epoch and −2 means the automatic value selected by the *h2o* package. The max number of times to iterate the whole dataset (epochs) is set as 500. The starting value of momentum is 0 or 0.5 (default 0, without hyperparameter grid search). The momentum damps the oscillation to achieve the optimal point and accelerates the iterations for faster convergence. The adaptive learning rate decay factor (rho) is 0.5 or 0.99 (default 0.99, without hyperparameter grid search). The quantile value (quantile_alpha value in *h2o*), when running quantile regression, is set between 0 and 1., Quantile regression is similar to linear regression, but measures the conditional quantile rather than the conditional mean of the response variable. The threshold between quadratic and linear loss (huber_alpha value in *h2o*) is set between 0 to 1 (default 0.9). “RandomDiscrete” strategy is used to enable search on all combinations of the hyperparameters. As part of the automatic machine-learning training, the maximum number of models for each run is set to 100. The training steps stop if the misclassification values do not improve by 0.01 after 5 iterations. Score_duty_cycle, the frequency of computing validation metrics, is set to 0.025 in *lilikoi v2.0*, meaning that no more than 2.5% of the total training time should be used to build the validation metrics.

For the exemplary ER dataset, after grid search, the final hyperparameters for its deep learning model are set as the following: “Rectifier” activation function, four 25-neuron hidden layers, input dropout ratio 0, default training samples per iteration per *h2o* (value of −2), epoch value of 430.9, momentum starting value 0, a rho value of 0.99, a quantile regression value of 1, and a huber alpha value of 0, other hyperparameters including an L1 regularization value of 2.5e-5, and an L2 regularization value of 2.6e-5.

This deep learning algorithm is added in classification along with other 6 machine learning techniques previously implemented in *Lilikoi v1*, namely generalized boosted model (GBM), linear discriminant analysis (LDA), logistic regression (LOG), random forest (RF), recursive partitioning and regression analysis (RPART), support vector machine (SVM). An n-fold cross validation (default n=10) is applied to avoid overfitting. Classification metrics such as Area Under the Curve (AUC), F1-statistic, balanced accuracy, sensitivity (SEN), and specificity (SPEC) are reported as bar plots.

### Prognosis prediction

*Lilikoi* v2.0 enables prognosis prediction, at either the metabolite level using metabolite-sample matrix, or the pathway-level (after *pathifier* based pathway transformation) using pathway deregulation score (PDS)-sample matrix. Currently two prognosis prediction methods are implemented: Cox-Proportional Hazards (Cox-PH) method (26) with penalization, and the neural-network based Cox-nnet method (5). *Cox-PH* is a survival regression model developed by David Cox in 1972. The input parameters are event (eg. death), survival time and penalized covariates: alpha to determine which penalization method to use, and lambda (lambda.min or lambda.1se) for prediction. Penalization is achieved by Lasso, Ridge or Elastic net with the *glmnet* package (27).

*Cox-nnet* is based on the artificial neural network (ANN) framework with a default of two-layer neural network: a hidden layer and an output layer (28). The output layer is fit to the Cox regression. *Lilikoi* v2.0 imports *Cox-nnet* originally written in Python, using the *reticulate* package.

The hazard function of the Cox-PH model is:

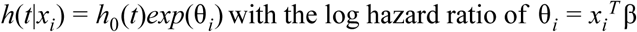

with its partial likelihood cost function :

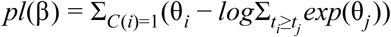

The Cox-nnet expands the Cox-PH hazard function above as:

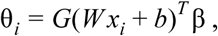

where *x*_*i*_ is the output of the hidden layer, G is the activation function and W is the coefficient weight matrix between the input and hidden layer.

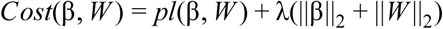

In the demonstration NCI data, we applied cross validation on the training dataset to determine the optimal L2 regularization lambda parameter. Cox-nnet supports two gradient descent algorithms, nesterov gradient descent and momentum gradient descent. Hyper-parameters can be set by users, including the gradient descent algorithm, initial learning rate, proportion of momentum, decrease of the learning rate, increase of the learning rate, number iterations between cost functions to determine increase or decrease of the learning rate, maximum number of iterations, stopping threshold, minimum number of iterations before stopping, number of iterations for new lowest cost before stopping and the random seed. The details can be found in the user manual.

The prognosis model is visualized by Kaplan-Meier curve plot, using the *survminer* package (29). Samples are dichotomized into different risk groups by prognosis index (PI), the logarithm of hazard ratio of the prognosis model. Lilikoi v2.0 allows several approaches for dichotomization: median PI threshold, event/non-event ratio, and quantile PI threshold (samples with PIs under the 1st quantile as the low-risk group and those above the 3rd quantile as the high-risk group).

The fitness of the models is evaluated by two metrics: C-index and log rank p-values. C-index is a goodness of fit measure of survival models (30). A C-index of 1 indicates that the model is the best model for prediction and C-index = 0.5 means that the model prediction is no better than a random guess. Log-rank p-value is based on the log-rank test (31, 32) to evaluate the null hypothesis that no difference in survival exists between the high-risk and low-risk groups. Log-rank p-value less than 0.05 means that there is significant difference between these two groups. Users have the option to split the data by N folds cross-validation, where the model is trained on the N-1 fold data and evaluated on the rest 1 fold data.

### Pathway level analysis

The selected pathway features from classification or prognosis prediction, can be visualized with the *Pathview* r package (33). Currently, any KEGG pathway can be used as the input to render pathway graphs. The top pathways are selected with the *featureSelection()* function in *Lilikoi*. Additionally, if there are corresponding gene expression profiles, they can be integrated with metabolites in *Pathview.*

The relationship between pathway and the metabolites in that particular pathway can be analyzed by single variate regression. The metabolites that are significantly associated with the pathway are displayed as bar graphs and top tables. All pathway features and their significantly associated metabolites are visualized by a bipartite graph with Cytoscape style. Cytoscape modules are imported in *Lilikoi* by the *RCy3* R package (34).

### Code availability

Lilikoi v2.0 source code with documentation and scripts to run testing data are available at https://github.com/lanagarmire/lilikoi2. Lilikoi v2.0 R package is submitted to CRAN team and upon passing, it will be expected to be available at: https://cran.r-project.org/web/packages/lilikoi/index.html.

## Results

### Overview of updated functionalities in *Lilikoi v2.0*

*Lilikoi v2.0* package is a significant upgrade of the previous version. It keeps all four modules in the original *Lilikoi v1* package: feature mapper, dimension transformer, feature selector, and classification predictor (2). However, given the recent applications of deep-learning in the metabolomics and other genomics fields (3–8), it is important to enable metabolomics researchers to investigate such new approaches. We thus implemented deep-learning as a new method in the classification module. Moreover, metabolomics have the potential to be prognosis markers (10), however, currently rarely metabolomics data analysis workflow is available to handle this issue. We herein implemented multiple methods for prognosis prediction, including *Cox-Proportional Hazard* (or *Cox-PH*) model and *Cox-nnet*, a neural-network based model (5). Additionally, we augmented the pathway-based metabolomics analysis with metabolite-pathway relationship analysis and pathway visualization. Last but not least, we also include additional preprocessing methods for metabolomics data analysis (eg. normalization, imputation) and tools for exploratory data analysis (eg. PCA and t-SNE analysis, and source of variation analysis).

Importantly, *Lilikoi v2.0* has added the following new functionalities (marked in red text boxes), as shown in Fig. 1. A pre-processing module is added for the initial steps, where normalization and imputation are considered. A new exploratory data analysis module is also added, to enable dimension reduction analysis (PCA or t-SNE) and source of variation analysis (SOV). The classification module is amended with the new deep-learning method, along with the previously implemented machine-learning methods. Additionally, a new prognosis module is introduced in this version, where cox-PH method and a new neural-network based Cox-nnet method are implemented. Downstream analysis and interpretation of pathways is also a new add-on feature, where visualization and metabolite-pathway regression are available.

**Figure 1:**
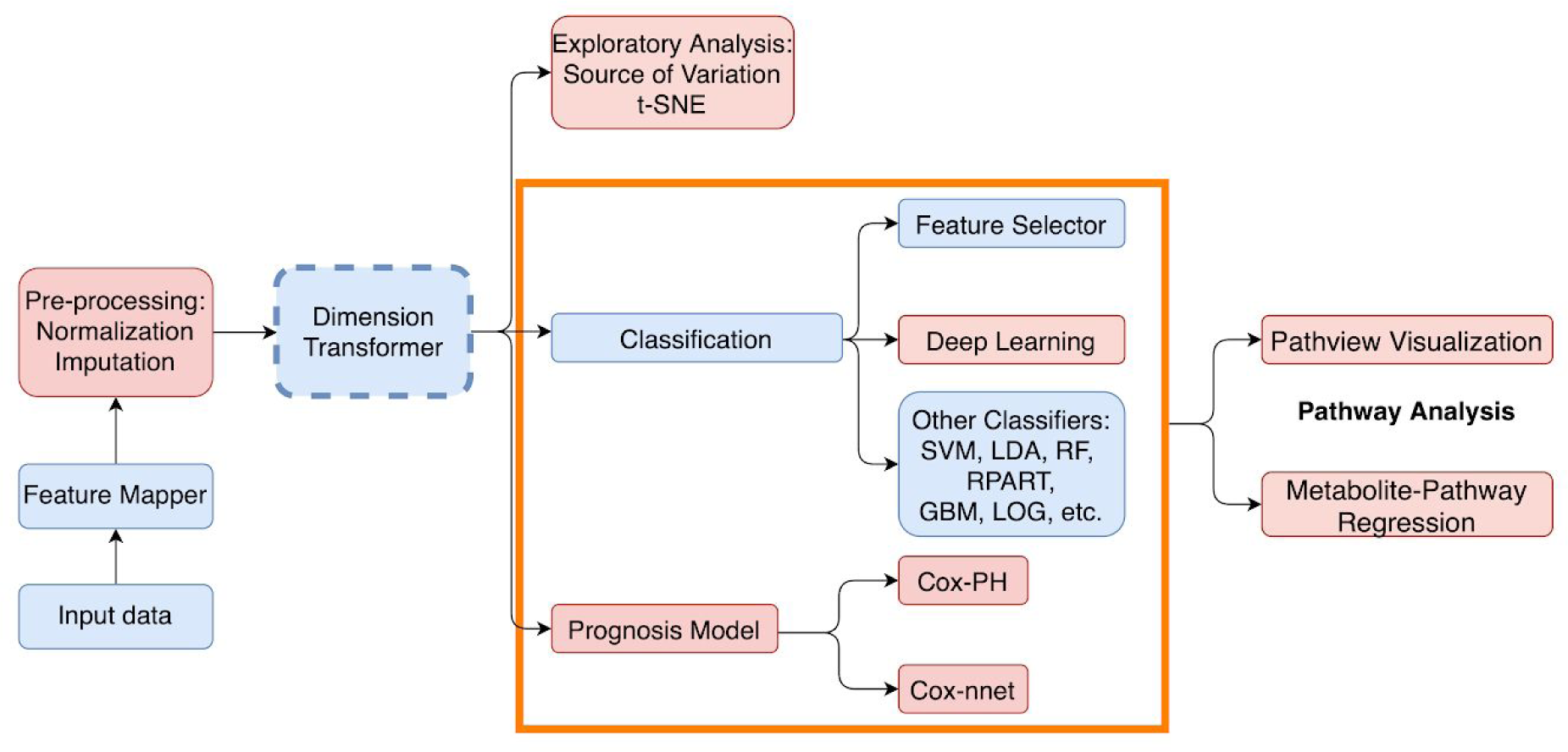
The workflow of *Lilikoi v2.0* package. *Lilikoi v2.0* is composed of seven modules: feature mapper, pre-processing, dimension transformer, exploratory analysis, classification, prognosis model, and pathway analysis. The pink boxes are new functionalities added to *Lilikoi v2.0*. Blue boxes are pre-existing modules in *Lilikoi v1.* Dashed box indicates an optional step.

### Data preprocessing and exploratory analysis

For data preprocessing, we added normalization and imputation methods. Three normalization methods (standard normalization, quantile normalization and median-fold normalization) are implemented, with median-fold normalization as the default method. For imputation of missing values, k-nearest neighbor method is the default method.

Un-supervised exploratory analysis is an important step to better understand the pattern in metabolomics data, as well as the metabolomics-phenotype relationship. To enable this, *Lilikoi* v2.0 added Principle Component Analysis (PCA) and t-Distributed Stochastic Neighbor Embedding (t-SNE) plot that help users to visualize high-dimensional metabolomics data. PCA reduces the dataset dimensions by finding out the linearly independent dimensions based on the eigenvalues and eigenvectors of the covariance matrix to represent the data. Different from the linear dimension reduction of PCA, t-SNE maps the high-dimensional data onto a low-dimensional space via a non-linear algorithm.

To investigate the metabolomics-phenotype data relationship, *Lilikoi* v2.0 has added the source of variation analysis between confounders and metabolomics data, based on ANOVA tests (18). Any clinical confounder with F-score bigger than the error term, whose F-score is 1, needs to be adjusted for in differential metabolite tests, when using other clinical variable(s) for grouping.

### Deep learning enabled classification module

Deep-learning enabled classification module is one of the highlighted functionalities of *Lilikoi v2.0*. The deep learning framework uses the same dataset and adopts the same architecture as previously described (9). The objective is to classify the 204 ER+ samples from the 67 ER-samples. We split the data roughly 4:1 ratio into training and testing data, with 10 fold cross-validation in the training data. We repeated this process 10 times randomly, to obtain averaged metrics.

We used the metabolite features as the inputs for deep-learning based classification, along with other popular methods: LDA, SVM, RF, RPART, LOG, and GBM (Methods). As shown in Fig. 2A and Table 1, deep learning on average performs the best overall in the training data, with the significantly higher F-1 statistic value (0.95) and sensitivity (0.98) than all other methods. F-1 statistic is a good unbiased metric given the unbalanced samples in ER= and ER-classes. However, the specificity (0.75) in the training dataset is second to the lowest (SPEC of LDA=0.72). The advantage of deep learning is more pronounced in the testing dataset (Fig. 2B and Table 1), where it achieves the highest values in AUC=0.91, SEN=0.95, and F1 statistics=0.93. Again the specificity is lower than other methods (0.69), probably due to the size of the samples.

**Table 1.** Performance of classification models on training and hold-out testing dataset.

| Data Set | Algorithm | AUC | SENS | SPEC | F1-Statistic | Balanced accuracy |
| --- | --- | --- | --- | --- | --- | --- |
| Training | DL | **0.909** | **0.978** | 0.747 | **0.952** | 0.777 |
|  | GBM | 0.906 | 0.600 | 0.945 | 0.666 | 0.772 |
|  | LDA | 0.700 | 0.583 | 0.718 | 0.478 | 0.651 |
|  | LOG | 0.906 | 0.608 | 0.946 | 0.681 | 0.777 |
|  | RF | 0.892 | 0.568 | 0.946 | 0.648 | 0.757 |
|  | RPART | 0.801 | 0.605 | 0.895 | 0.620 | 0.750 |
|  | SVM | 0.905 | 0.663 | 0.920 | 0.688 | 0.791 |
| Testing | DL | **0.912** | **0.954** | 0.688 | **0.930** | 0.747 |
|  | GBM | 0.878 | 0.560 | 0.939 | 0.639 | 0.749 |
|  | LDA | 0.745 | 0.627 | 0.754 | 0.527 | 0.691 |
|  | LOG | 0.873 | 0.550 | 0.943 | 0.634 | 0.747 |
|  | RF | 0.870 | 0.578 | 0.938 | 0.643 | 0.758 |
|  | RPART | 0.767 | 0.609 | 0.861 | 0.589 | 0.735 |
|  | SVM | 0.883 | 0.653 | 0.927 | 0.693 | 0.790 |

**Figure 2:**
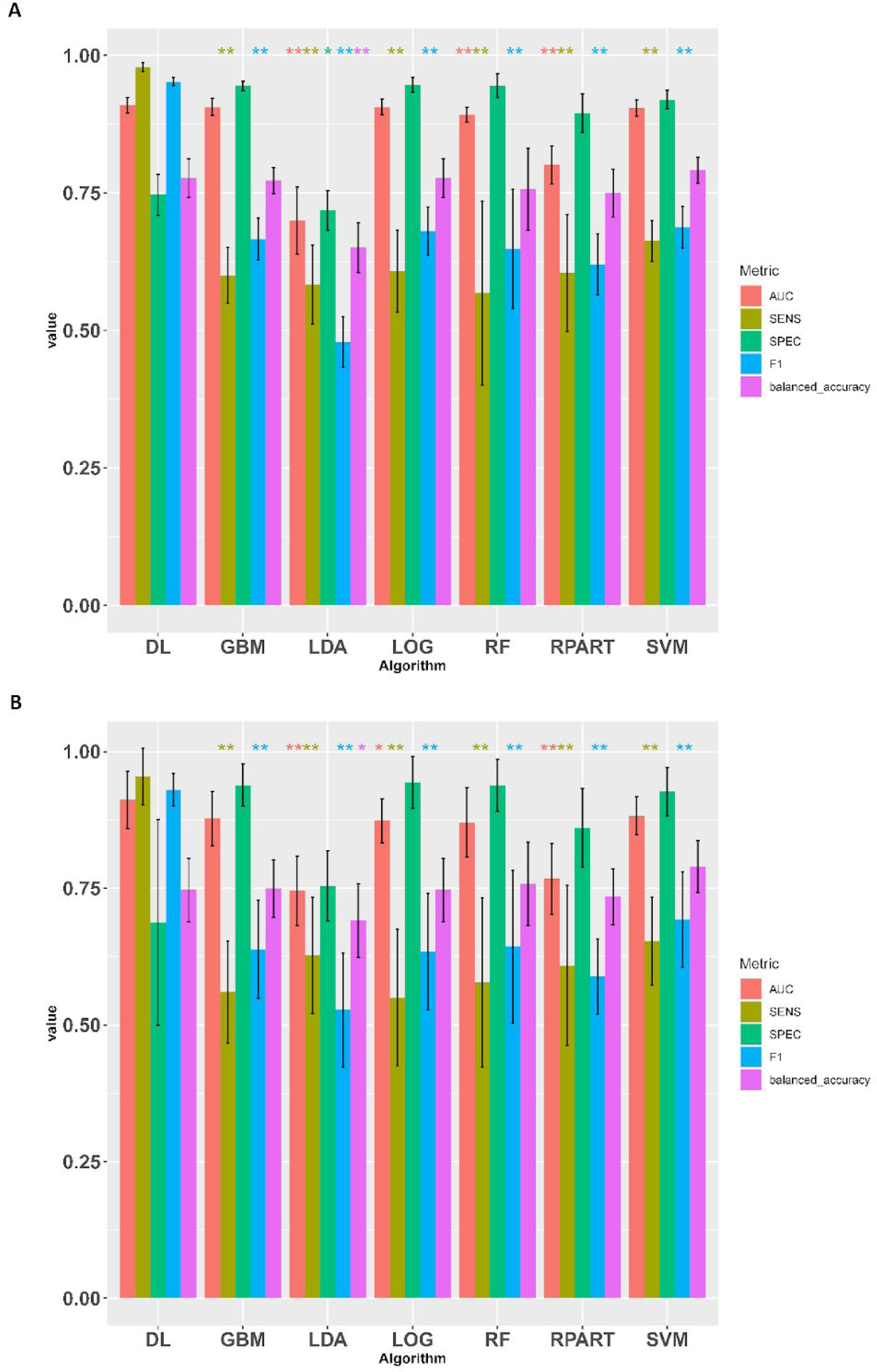
Model evaluation on deep learning (DL) and other machine learning techniques. (A) metrics on training data sets; (B) metrics on testing data sets. Abbreviations: deeplearning (DL), generalized boosted model (GBM), linear discriminante analysis (LDA), logistic regression (LOG), random forest (RF), recursive partitioning and regression analysis (RPART), support vector machine (SVM). *: p-value < 0.05 (one-tail t-test) compared to the same metric in DL; **:p-value < 0.01;

### Prognosis prediction

Deep-learning enabled prognosis prediction is another unique functionalities of *Lilikoi v2.0*, compared to other metabolomics analysis packages and toolkits. To demonstrate prognosis analysis, we used the NCI dataset as described in Methods. As the unique feature of Lilikoi is pathway-level modeling, the metabolites intensity data are first transformed to pathway level data matrix (see Methods). Penalized survival analysis using *Cox-PH* model and *Cox-nnet* were conducted. For *Cox-PH* regression, L2 norm (Ridge) penalization was applied to select featured pathways. After fitting, the prognosis index (PI) was used to separate the patient into the high-risk vs low-risk groups using the first quantile of PI as the threshold. As shown by the Kaplan-Meier curves in Fig. 3, the *Cox-PH* model yields a C-index value of 0.65 and log rank p-value of 0.04 (Fig. 3A); *Cox-nnet* model yields slightly better results, with a C-index value of 0.66 and log rank p-value of 0.02 (Fig. 3B).

**Figure 3:**
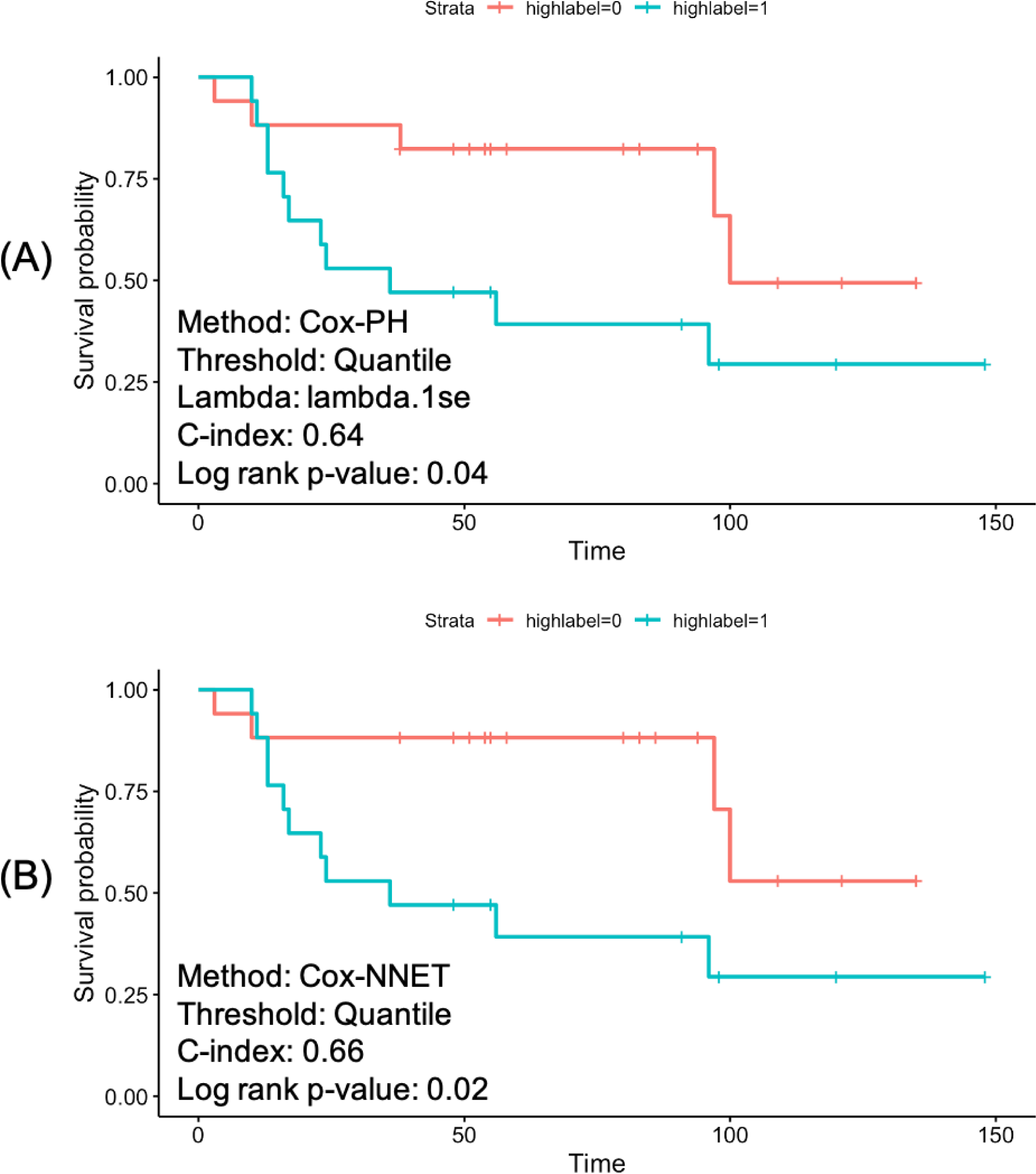
Comparison of Kaplan-Meier curves resulting from Cox-PH and Cox-nnet. The samples are dichotomized into 2 risk groups by the first quantile of the prognosis index (PI) score. (A) Cox-PH model. (B) Cox-nnet model with 3-layer neural network: one input layer, one fully connected hidden layer and the output layer.

### Pathway downstream analysis

We used the metabolites expression information in the aforementioned workbench breast cancer dataset PR000284 as the cpd.data input of the *pathview* function. According to our *featureSelection* results, alanine aspartate and glutamate metabolism is one of the top pathways for metabolite data. Therefore, we demonstrate the pathway visualization, based on the *Pathview* R package. using “alanine aspartate and glutamate metabolism pathway” (Fig. 4). As shown in Fig. 4, 6 metabolites in this pathway have intensities. Asparagine has increased levels in ER-patients, due to the conversion from its substrate aspartate, which is reduced in ER-patients. The reduction of aspartate in ER-patients is consistent with observation before (35).

**Figure 4:**
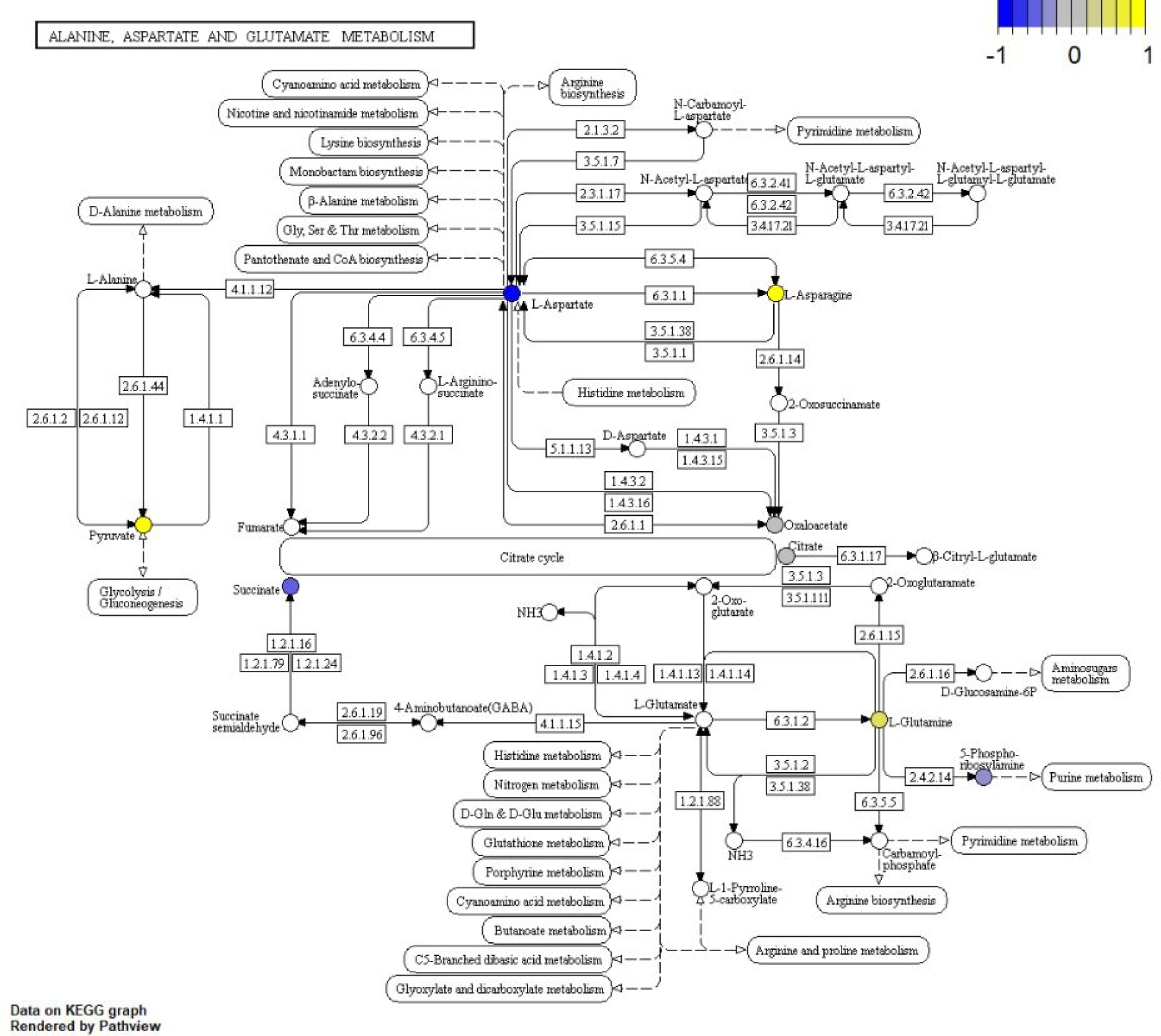
Pathway visualization: alanine aspartate and glutamate metabolism pathway. Color scheme is based on log2 transformed ratio of the mean values of ER-samples over ER+ samples. The pathway rendering is done by the *Pathview* R package.

It is important to link the significant metabolites that contribute to the pathway features. For this, single-variate regressions between metabolites and pathways are conducted, with the workbench dataset with 207 plasma samples (126 breast cancer cases and 81 control cases). The regression results (Fig. 5) can be visualized by the partite graph, where the yellow nodes represent pathway features, and the green nodes are metabolites significantly (p<0.05) associated with the pathways. show how each metabolite contributes to the selected pathways. The generic term “metabolic pathways”, is associated with the largest number (86) of metabolites. Among them, isopentenyl pyrophosphate has the most weight on the edge. Many pathways related to amino acid synthesis and metabolism are highlighted. Users can also elect to exam the metabolites within a particular pathway, by individual bar graphs. As an example, we show the metabolites that are associated with “alanine aspartate and glutamate metabolism pathway” (Fig. 5, insert). Citric acid, pyruvate, 5-phosphoribosylamine, glutamine, oxaloacetate and asparagine all significantly (p<0.05) increase in ER-patients, with coefficients of 0.043, 0.046, 0.049, 0.378, 0.575 and 0.997 from single-variate linear regressions; on the other hand, succinate and aspartate have opposite significant decreases, with coefficients of −0.435 and −0.269. Additional bar graphs showing relationships of metabolites and all top 10 pathways are in Supplementary Figure S1.

**Figure 5:**
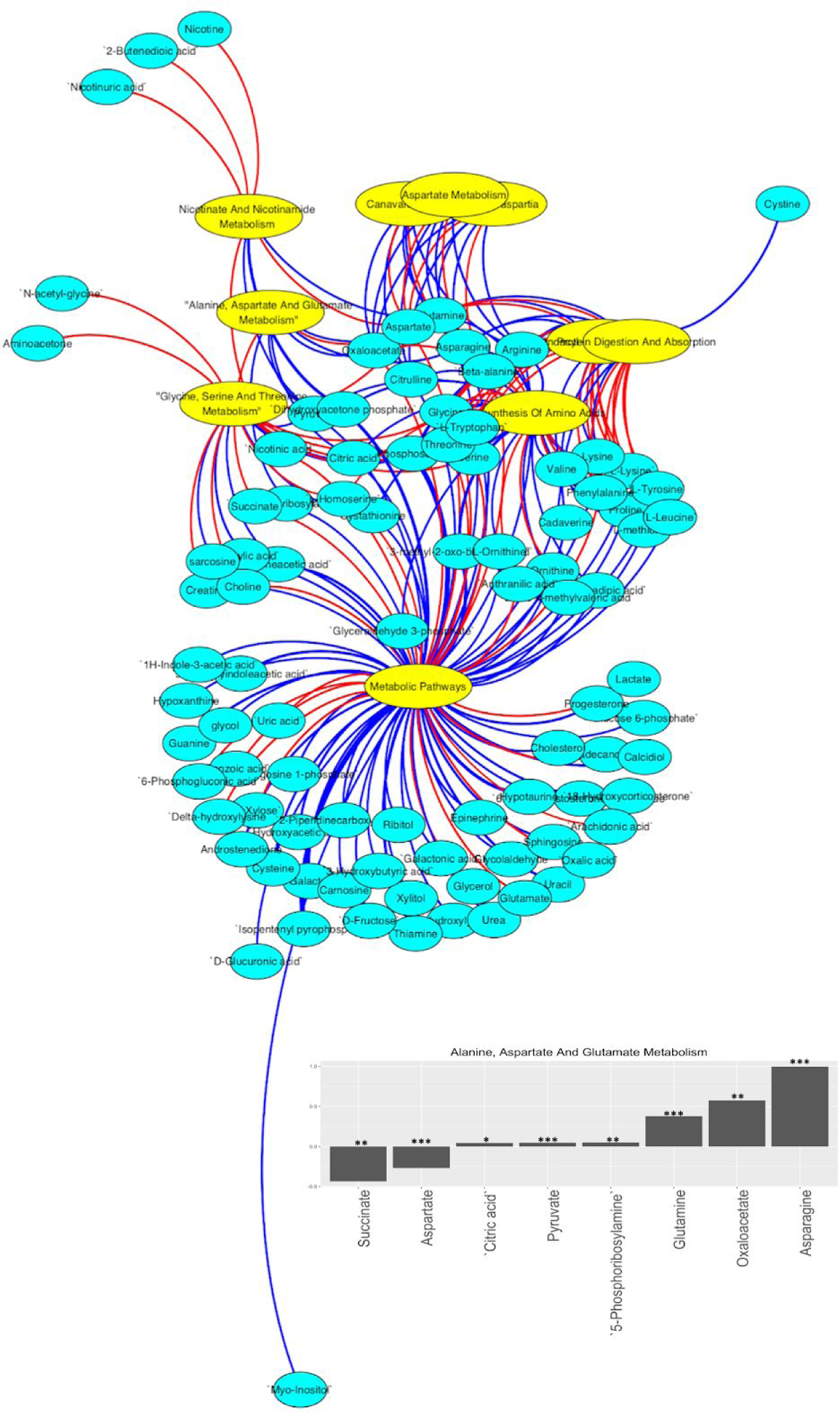
Metabolite-pathway relationship analysis. Bipartite plot with top 10 pathways and corresponding metabolites. Cyan and yellow nodes indicate metabolites and pathways, respectively. Red and blue edges are negative (-) and positive (+) associations, respectively. Thicker edges indicate higher levels of association. Insert: Bar plots of the relationship between the Alanine, Aspartate And Glutamate Metabolism pathway and its corresponding metabolites. *: p<0.05; **: p<0.01; ***:p<0.001.

## CONCLUSIONS

Here we report the upgrade of *Lilikoi* v2.0, a new deep-learning enabled, personalized pathway-based package for diagnosis and prognosis predictions using metabolomics data. The new version of *Lilikoi* added many new modules, ranging from data preprocessing, exploratory analysis, deep learning, prognosis prediction, and visualization. Building upon the previous work on pathway-based modeling and prediction, *Lilikoi* v2.0 allows much better exploration of pathway-based analysis using various modern analytics methods for classification and survival analysis, including deep learning implementation. Such endeavor sets Lilikoi apart from other more conventional metabolomics analysis packages (36–38).

Some practical challenges still exist, leaving room for the future development of *Lilikoi*. For example, mapping rate of metabolites and pathways can be further improved, by using better matching algorithms and/or literature mining with Natural Language Processing (NLP). Also, the current best classification model in *Lilikoi* is determined by users. We would like to automatically recommend the best classification model for users. Implementing automatic machine learning algorithms (AutoML), as suggested by Auto-WEKA (39) and other applications (40), will be considered for future classification modules. Moreover, integration between metabolomics and other genomics data types is becoming increasingly important, and will be modeled in the next version of *Lilikoi*, potentially with deep-learning and machine-learning ensemble tools, such as DeepProg models that are developed by us and others (3, 4, 8, 41).

## ACKNOWLEDGEMENTS

This research was supported by grants K01ES025434 awarded by NIEHS through funds provided by the trans-NIH Big Data to Knowledge (BD2K) initiative (www.bd2k.nih.gov), R01 LM012373 and R01 LM012907 awarded by NLM, and R01 HD084633 awarded by NICHD to L.X. Garmire.

## Notes

### Competing Interest Statement

The authors have declared no competing interest.

