## Supplement Figure 1 for "Lilikoi V2.0: a deep-learning enabled, personalized pathway-based R package for diagnosis and prognosis predictions using metabolomics data"

### SUPPLEMENTARY FIGURE

**Figure S1: relationships between metabolites and all ten pathways.** Pathways are selected by the featureSelection function in *lilikoi,* with selection threshold of 0.54 and decision tree method of gain ratio.

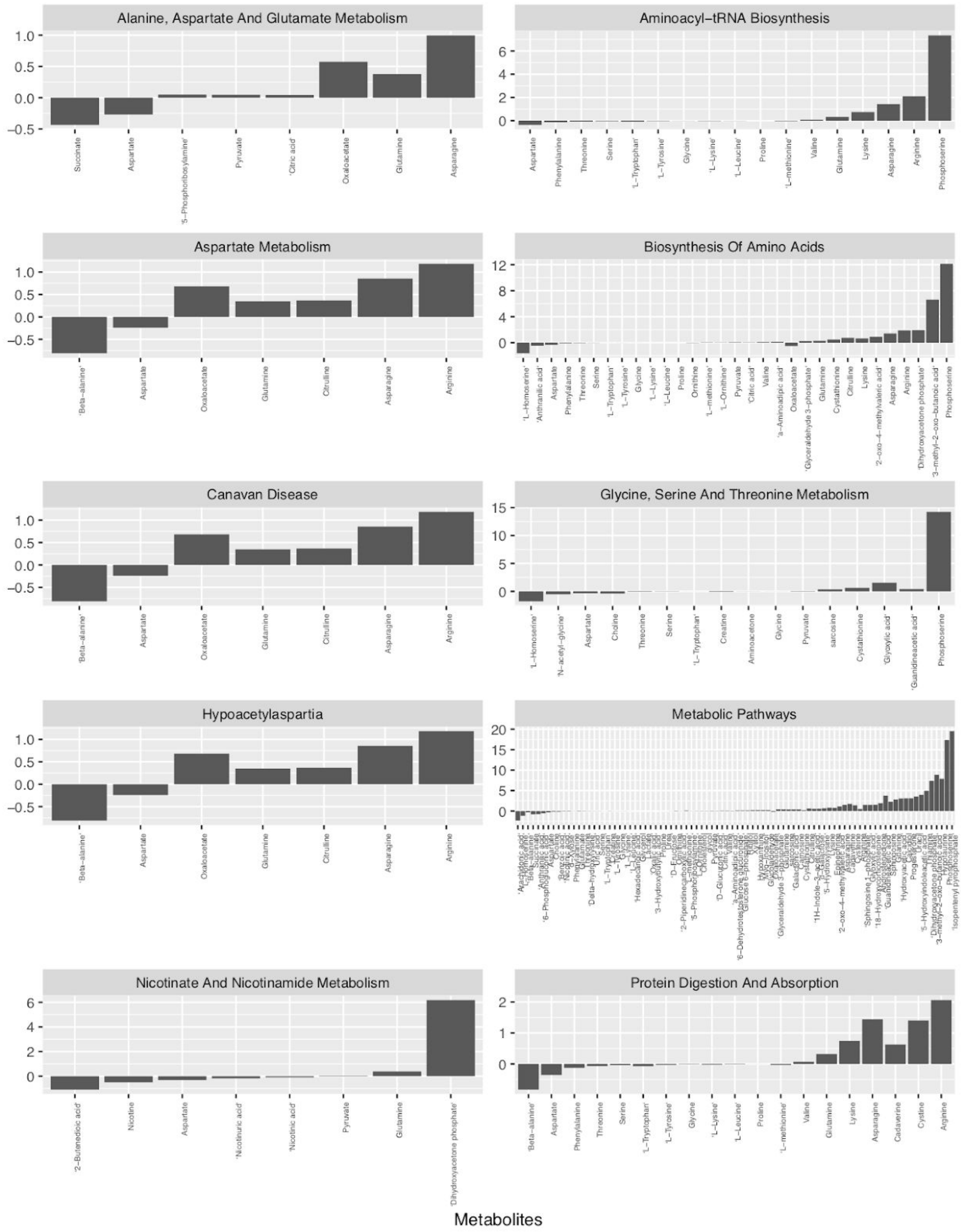
